# Dynamics of deep water and N uptake under varied N and water supply

**DOI:** 10.1101/2021.09.27.461951

**Authors:** Guanying Chen, Camilla Ruø Rasmussen, Dorte Bodin Dresbøll, Abraham George Smith, Kristian Thorup-Kristensen

## Abstract

**Aims:** Enhanced nitrogen (N) and water uptake from deep soil layers may increase resource use efficiency whilst maintaining yield under stressed conditions. Winter oilseed rape (*Brassica napus* L.) can develop deep roots and access deep-stored resources such as N and water, while this potential has large uncertainties in variable environments. In this study, we aimed to evaluate the effects of reduced N and water supply on deep N and water uptake.

**Methods:** Oilseed rape plants grown in outdoor rhizotrons were supplied with 240 and 80 kg N ha^-1^ respectively in 2019 whereas a well-watered and a water-deficit treatment were established in 2020. To track deep water and N uptake, a mixture of ^2^H_2_O and Ca(^15^NO_3_)_2_ was injected into the soil column at 0.5 and 1.7 m depths. δ^2^H in transpiration water and δ^15^N in leaves were measured after injection. δ^15^N in biomass samples were also measured.

**Results:** Differences in N or water supply had little effect on root growth. The low N treatment reduced water uptake throughout the soil profile, but caused a non-significant increment in ^15^N uptake efficiency at both 0.5 and 1.7 m. Water deficit in the upper soil layers led to compensatory deep water, while N uptake was not altered by soil water status.

**Conclusion:** Our findings demonstrate that for winter oilseed rape, high N application and water deficiency in shallow layers increases deep water uptake, and that the efficiency of deep N uptake is mainly sensitive to N supply rather than water supply.

## Introduction

Nitrogen (N) and water are the main factors that can be modified in agricultural production and have been widely documented for their crucial roles in determining yield (Mueller et al. 2012; Sinclair and Rufty 2012). Inadequate supply of N and water leads to yield loss, while the loss of N from agricultural land also causes environmental problems. Therefore, improving water and N use is crucial for sustainable agricultural production.

Deep-rooted crops have shown great potential in enhancing soil N and water uptake, as well as improving yield (Wasson et al. 2012; Thorup-Kristensen et al. 2020a), while deep root development and function is highly sensitive to the external environment (Lynch and Wojciechowski 2015). Management and environmental factors such as N and water supply affect plant growth, soil N and water availability, and subsequently affect root uptake. N supply affects overall plant growth and yield (Asare and Scarisbrick 1995; Khan et al. 2017), but the current findings of the effects of N supply on root growth and N uptake seem ambiguous. Increasing N supply stimulates root growth either via more robust shoot growth or as a consequence of increased soil N availability (Hodge et al. 1999). In contrast, Svoboda and Haberle (2006) found that the rooting depth and deep root density of winter wheat (*Triticum aestivum* L.) could be reduced when increasing N fertilization. Although plant root surface area and total N uptake may be enhanced by a higher fertilization rate (Lynch et al. 2012), it has been reported that nitrogen uptake efficiency in oilseed rape (*Brassica napus* L.) and winter wheat can decline with increasing N fertilizer rate (Rathke et al. 2006; Rasmussen et al. 2015). Riar et al. (2020) found that, independent of irrigation, oilseed rape plants fertilized with 100 kg N ha^−1^ had higher N uptake efficiency than those fertilized with 200 kg N ha^−1^.

Nitrogen supply is important for improving water use. Increased N application may result in higher water use efficiency of wheat and oilseed rape (Taylor et al. 1991; Waraich et al. 2011), possibly due to improved shoot growth. Compared with no or less fertilization, adequately fertilized crops usually grow more vigorously and have larger leaf areas, increasing transpiration and decreasing soil evaporation. Increasing N supply increases transpiration intensity under normal water supply, thus enhancing water use (Li et al. 2009).

Water availability controls crop growth, especially canopy development, but it also controls root growth. Bloom et al. (1985) hypothesized that plant root growth may be stimulated under drought to enhance or maintain the capacity for acquiring water. Accordingly, Vandoorne et al. (2012) found that although chicory’s total root length decreased under water deficit conditions, the overall root profiles developed deeper and triggered compensation from wetter and deeper horizons. Furthermore, both Li et al. (2011) and Álvarez et al. (2011) observed a significant increase in water use efficiency during water deficit.

In addition to altering the root growth and water acquisition, water status affects plant N uptake in various ways. Under the same fertilization rate, Riar et al. (2020) showed that irrigation management enhanced soil moisture and further improved oilseed rape’s N uptake efficiency and N use efficiency by 40%. Water deficit also affects nitrogen demand, nitrogen availability and nitrogen assimilation and partitioning of the assimilates (Sadras et al. 2016). The reduced shoot growth driven by water deficit reduces plant nitrogen demand and tends to decrease nitrogen use efficiency if N input are not reduced correspondingly (Quemada and Gabriel 2016). The availability and supply of soil N can be limited by soil dryness due to reduced soil organic N mineralization (Jensen et al. 1997) and restricted nitrate movement by both mass flow and diffusion (Plett et al. 2020). Water deficit could also diminish nitrate reductase activity, hence reducing plant N assimilation (Gonzalez-Dugo et al. 2010). Moreover, assimilates tend to translocate to the roots rather than the shoots in the case of water deficiency (Li et al. 2011). In summary, both N and water supply can affect root growth, water, and N use.

Deep roots are not assumed to be as efficient as shallow roots in N and water uptake, as the roots reach the subsoil layers late in the growing period and are not able to develop as high densities there as in the topsoil. Existing studies indicate that water deficiency in topsoil can increase deep water uptake (Kirkegaard et al. 2007) and that decreased N supply in topsoil increased deep nitrogen uptake (Kuhlmann et al. 1989; Haberle et al. 2006). However, it is less clear to what extent deep root growth, as well as deep uptake of N and water, are affected by the total N and water availability for a crop. Studies of deep root growth, water, and N use under different N and water regimes could increase our understanding of the functions of deep roots. In addition, such studies would increase the understanding of the contribution of deep roots to crop N and water supply and to reduce N leaching losses, and how this is affected by crop management.

Oilseed rape is known for its high capacity for N and water uptake and has the potential to develop roots in soil layers below 2 m (Dresbøll et al. 2016; Kirkegaard et al. 2021). In this study, we used oilseed rape as the model crop and examined how N and water supply affect the root growth, utilization of N and water from deep soil layers, and N and water uptake dynamics in the subsoil. It was hypothesized that (I) N and water deficiency in topsoil stimulate root growth in deeper soil layers; (II) Lower N availability in the upper soil layer reduces total water uptake, but enhances N uptake from the subsoil. (III) Lower water availability in the upper soil layer reduces the N uptake from the whole soil profile, but increases water uptake from the subsoil.

## Materials and methods

### Experimental facility

Two consecutive experiments were conducted in the seasons 2018/2019 and 2019/2020 using the rhizobox facility (Thorup-Kristensen et al. 2020b) at the University of Copenhagen in Taastrup, Zealand, Denmark (55°40′ N; 12°18′ E). The facility consists of rhizoboxes that allow observations of root growth and root activity down to 4 m depth. The growth medium was field soil. Both years the topsoil was replaced right before planting (Table 1). The rhizoboxes are rectangular columns of 1.2 × 0.6 m, divided into an east- and a west-facing chamber, each with a surface area of 1.2 × 0.3 m. The front of the chambers is divided into 20 panels by metal frames covered by removable white foamed PVC boards, allowing root observations through transparent acrylic boards. The acrylic boards can be removed for sampling and measurements that require direct soil contact. For further details on the facility, see Rasmussen et al. (2020) and Thorup-Kristensen et al. (2020b).

**Table 1.** Characteristics of the soil in rhizoboxes

| Depth | Coarse sand (%)<br>0.2–2.0 mm | Fine sand (%)<br>0.02–0.2 mm | Silt (%)<br>0.002–0.02 mm | Clay (%)<br><0.002 mm | Organic matter<br>(%) |
| --- | --- | --- | --- | --- | --- |
| 0 - 0.2 m | 46.4 | 39.7 | 5.5 | 7.0 | 1.4 |
| 0.2 – 2.0 m | 26.8 | 51.1 | 4.2 | 17.6 | 0.3 |
| 2.0 – 4.0 m | 21.1 | 54.1 | 8.7 | 16.0 | < 0.1 |

### Experimental design

Oilseed rape (*Brassica napus* L., cv. “Butterfly”) plants were sown in the field on August 16, 2018 (Exp. 1) and in pots on August 13, 2019 (Exp. 2) before being transplanted to the rhizoboxes on October 8, 2018 and August 26, 2019, respectively. Due to a pest infestation *(Delia radicum)* in September 2019, a few plants were replaced by spare ones on September 24, 2019. The re-transplanted plants were smaller than the original ones during the entire growing period. Plant density in both years was five plants per chamber, corresponding to 14 plants m^-2^.

In Exp. 1, two N treatments were established by fertilizing with a nutrient solution, applying nutrients in a high N treatment (N240) equivalent 240 kg N ha^−1^, 38 kg P ha^-1^, 192 kg K ha^-1^; and a low N treatment (N80) equivalent to 80 kg N ha^−1^, 13 kg P ha^-1^, 65 kg K ha^-1^ respectively on March 27, 2019. During this season, all chambers received water through precipitation and irrigation, which were sufficient to keep them well watered. In Exp. 2, two irrigation regimes were established. Rainout shelters were mounted on top of all chambers on February 26, 2020, to allow complete control of soil moisture by irrigation. Well-watered (WW) chambers were irrigated with 60 mm water on April 14 and again on April 15, 2020 to establish soil profiles with high initial water content. No more irrigation was given to the well-watered chambers until May 10, 2020. In the following month, the well-watered chambers were irrigated frequently to keep an adequate water supply. Water deficit (WD) chambers received no irrigation during the whole experimental period. In Exp. 2, all chambers were fertilized with a total of 200 kg N ha^−1^, 38 kg P ha^-1^, 192 kg K ha^-1^. Fertilization was divided into three applications, with N supply of 40, 80, and 80 kg N ha^−1^ on September 5, 2019, March 2, and April 1, 2020. The treatments and timeline of the experiments are shown in Table 2. The two treatments in Exp. 1 and 2 were established in six randomly distributed replicates.

**Table 2.** Treatments and timeline of the experiments. The oilseed rape plants were fertilized with 240 kg N ha^-1^ (N240) or 80 kg N ha^-1^ (N80) applications in Exp. 1; in Exp. 2 plants were grown under well-watered (WW) or water deficient (WD) conditions at the beginning of the labelling.

|  | Exp. 1<br>2018/2019 | Exp. 2<br>2019/2020 |
| --- | --- | --- |
| Sowing | 16 August 2018 | 13 August 2019 |
| Transplanting | 8 October 2018 | 26 August 2019 <sup>a</sup> |
| Rate of N application | N240: 240 kg N ha<sup>-1</sup><br>N80: 80 kg N ha<sup>-1</sup> | 200 kg N ha<sup>-1</sup> |
| Date of N application | 27 March | 1<sup>st</sup>: 5 September 2019<br>2<sup>nd</sup>: 2 March<br>3<sup>rd</sup>: 1 April |
| Water status | WW: Well-watered | WW: Well-watered<br>WD: Water deficit |
| Key irrigation events <sup>b</sup> | None | WW 1<sup>st</sup>: 14 April, 20 mm<br>WW 2<sup>nd</sup>: 15 April, 40 mm<br>WD: None |
| Tracer injection | 3 April | 17 April |
| Isotope sampling | 1<sup>st</sup>: 2 April<br>2<sup>nd</sup>: 10 April<br>3<sup>rd</sup>: 17 April<br>4<sup>th</sup>: 23 April<br>5<sup>th</sup>: 1 May | 1<sup>st</sup>: 16 April<br>2<sup>nd</sup>: 22 April<br>3<sup>rd</sup>: 27 April<br>4<sup>th</sup>: 3 May<br>5<sup>th</sup>: 8 May |
| Final biomass collection | 5 June | 18 June |

<sup>a</sup> Pest infestation (*Delia radicum*) occurred in some of the chambers in September 2019. Thus, a few plants were replaced on September 24, 2019. Replaced plants were sown together with the original ones, but appeared smaller than the original ones throughout the experiment. <sup>b</sup> Irrigation events which might change the soil water status before tracer injection were counted.

### ^2^H and ^15^N labelling

In both experiments, water and N uptake were traced using isotope labelled water and N injected into the soil at either 0.5 or 1.7 m depths. Tracer application was repeated in three chambers for each depth and treatment. Tracers were injected when the roots had already reached 1.7 m depth. The tracer application rates aimed at ensuring significant enrichment in plants and transpiration water, and were based on estimated N and water availability in the soil, the natural isotope enrichment, and assumed uptake rates of applied tracers. In Exp.1 where two N fertilizer levels were established (240/80 kg ha^-1^), the ^15^N application was adjusted similarly and the N240 and N80 treatments received 0.96 g and 0.32 g ^15^N, respectively. In Exp. 2, each chamber received 0.5 g ^15^N. Tracer solution was prepared by mixing the specific amount of Ca(^15^NO_3_)_2_ (>98.9 at% ^15^N) with 50 ml ^2^H_2_O (^2^H content = 99.94%) and 50 ml distilled water for each chamber.

The tracer was injected into 20 injection holes at each injection depth, which were evenly distributed in two parallel rows. The holes were 25 cm deep, made by a steel stick 0.5 cm in diameter. Inside each of the 20 holes, a 5 ml tracer solution was injected. The syringe needle was pushed 25 cm into the soil, and 1 ml of the solution was released every five centimeters as the syringe was drawn back. In this way, the tracer solution was distributed into 100 individual points in the soil at each injection depth. The injection procedures were conducted between 1:00 – 4:00 pm on April 3, 2019 and April 17, 2020.

### Sampling and sample preparation

Transpiration water for ^2^H tracing and leaf samples for ^15^N tracing was collected five times in each experiment. The first sampling time was in the morning, right before the injection, and subsequently four times after the injection (Table 2). The collection of transpiration water was initiated between 10:00 and 11:00 on the sampling day. Each plant was covered with a plastic bag that was tightened by a rubber band at the bottom. After two hours, the condensed droplets of transpired water inside the bags were collected. The water was quickly transferred from the bags to sealed plastic bottles. The collected transpiration water was filtered through 2 μm filter paper to remove dirt and debris. Filtered water from all plants grown in the same chamber was mixed for ^2^H analysis. Three to five of the latest fully developed leaves were collected on the same days as the transpiration water samplings. Leaf samples were dried, weighed, milled, and then encapsulated for ^15^N analysis. To determine the effect of water deficit on plant growth, the leaf samples were also analyzed for ^13^C in Exp. 2.

The total aboveground biomass was collected on June 5, 2019 and June 18, 2020, in Exp. 1 and 2, respectively. Biomass samples were divided into stems, pods, and leaves. However, in Exp. 2, all leaves had been shed when the total biomass was collected. Biomass samples from all plants in each chamber were mixed, dried at 70°C to constant weight, and weighed and stored until further analysis. In both experiments, biomass samples were analyzed for ^15^N.

Soil samples from 0.5, 1.1, and 1.7 m soil depths were taken before tracer injection and after the last isotope sampling to determine soil nitrate and ^15^N concentration. All soil samples were frozen immediately after sampling and stored until further preparation. Subsequently, 20 g soil was taken from each sample and mixed with 100 ml 2M KCl solution. The mixture was shaken for one hour and filtered through 2 μm filter paper. All solution samples were frozen for later analysis.

### Isotopic analyses

All isotopic measurements were done by the Stable Isotope Facility, UC Davis. ^15^N and ^13^C values in biomass samples were analyzed using IRMS. ^2^H values in transpiration water samples were analyzed using the Laser Water Isotope Analyzer V2 (Los Gatos Research, Inc., Mountain View, CA, USA). ^15^N concentration in soil samples was measured using IRMS. Nitrate-N content in the frozen soil solution was measured using the flow injection analyzer method.

^2^H and ^15^N enrichment (‰) was calculated as the increase of ^2^H and ^15^N values from pre-tracer sampling to post-tracer sampling unless otherwise stated. The ratio (%) of 1.7 m - and 0.5 m - derived ^2^H enrichment in transpiration water in the same treatment was calculated to investigate the distribution of water uptake. To compare ^15^N uptake between different treatments more directly, ^15^N uptake efficiency (^15^N_upe;_ % g ^-1^) was calculated as:

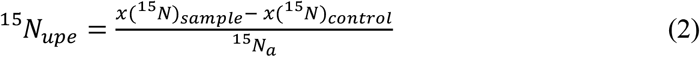

where *x*(^15^N)_sample_ and *x*(^15^N)_control_ are the atom fraction of ^15^N in post-tracer samples and pre-tracer samples, respectively. In harvest samples, *x*(^15^N)_control_ refers to the natural abundance of ^15^N in plant organs, which is usually 0.366%. ^15^N_a_ is the total amount (g) of ^15^N that was added to the soil.

### Soil water measurements

Four time-domain reflectometry sensors (TDR-315/TDR-315L, Acclima Inc., Meridian, Idaho) were installed in every chamber. They were placed at 0.5, 1.4, 2.3, and 3.5 m depth, respectively, recording soil volumetric water content (VWC; %) at least every 30 minutes. The VWC sensor readings were calibrated against VWC in soil samples taken in close proximity to the sensors. The samples were taken using metal rings with a diameter and a height of 5 cm and VWC was calculated based on the fresh and dry weight of the samples. For each sensor at least 3 samples were collected at different times aiming a covering a broad range of water content. Based on the correlations between sensor VWC and sample VWC, the sensor readings were adjusted to obtain an intercept of zero. The correlations did not call for a slope adjustment.

Two periods around the middle of the isotope sampling period were selected for estimating the water uptake during the sampling period. In Exp. 1, it was a 20-day period starting from 4 days after injection and ending four days before the last sampling date. In Exp. 2, a 14-day period was selected, which began four days after injection and ended four days before the last sampling date. Letting each sensor represent a 1 m depth-interval the soil water content in each interval was calculated (mm m^-1^ soil column). No water was added during the selected periods, thus soil water movement was assumed negligible and a decrease in soil water content was interpreted as plant water uptake. In both experiments, the daily water uptake was calculated only for the top 3 m soil columns, where most roots were found.

### Root imaging, segmentation, and calculation

During the experimental periods, the growth of oilseed rape roots was recorded every three to four weeks with a digital camera (Olympus Tough TG 860). The camera was in a box excluding daylight but with internal LED light strips as the light source. The box fits the frames of each panel of the rhizobox chambers, and by taking five photos per panel, the total area of each panel was photographed for subsequent image segmentation. RootPainter (Smith et al. 2020a) was used to segment roots from the soil background. A model trained with randomly selected images was used to segment roots on all the images and estimate the root length in each image via skeletonization and pixel counting (Smith et al. 2020b). Root intensity was calculated as cm of root per cm^2^ of soil in the images.

### Statistics

Data analyses were conducted in R (Version 3.5.3, R Core team 2019). The effect of N and water treatment on the harvested biomass and N content was tested in t-tests in separate tests for each experiment and for each plant organ. T-tests were used for comparing root intensity under different treatments. Separate tests were performed for each experiment and depth. The main effects of N/water supply and depth on daily water uptake were tested using a linear mixed model. A linear mixed model was used to examine differences in δ^13^C in leaf samples under different water treatments in Exp. 2. Foliar δ^13^C values from all sampling dates were compared together, where dates and water treatments were fixed effects and chamber was a random effect. Linear mixed models were used to examine differences in ^2^H enrichment in water samples and ^15^N_upe_ in biomass samples among N/water treatments, dates, and injection depths, where the combined factor of N/water level and depth (level-depth combined treatment, e.g., N80 - 0.5 m) and dates were fixed effects and chamber was a random effect. Multiple comparisons were conducted subsequently to test for changes in ^2^H enrichment and ^15^N_upe_ within the same level-depth combined treatment among all the dates. In both experiments, the effect of N/water supply on ^15^N_upe_ in harvest samples within the same organ was tested using linear mixed models with level-depth combined treatment as a fixed factor and chamber direction as a random factor. Multiple comparisons were done to compare ^15^N_upe_ in harvest samples within the same organ.

For ^2^H enrichment analysis, data were log-transformed to fulfill assumptions of normality and homogeneity. Multiple comparisons (Tukey HSD; P≤0.05) were based on values derived from linear mixed models.

## Results

### Biomass

Oilseed rape plants grew well in both years. In Exp. 1, the effect of N fertilization rate was evident, as the N240 treatment resulted in significantly higher leaf, stem, and pod biomass than N80 (Table 3). The N content in all three organs also increased when more N was given.

**Table 3.**
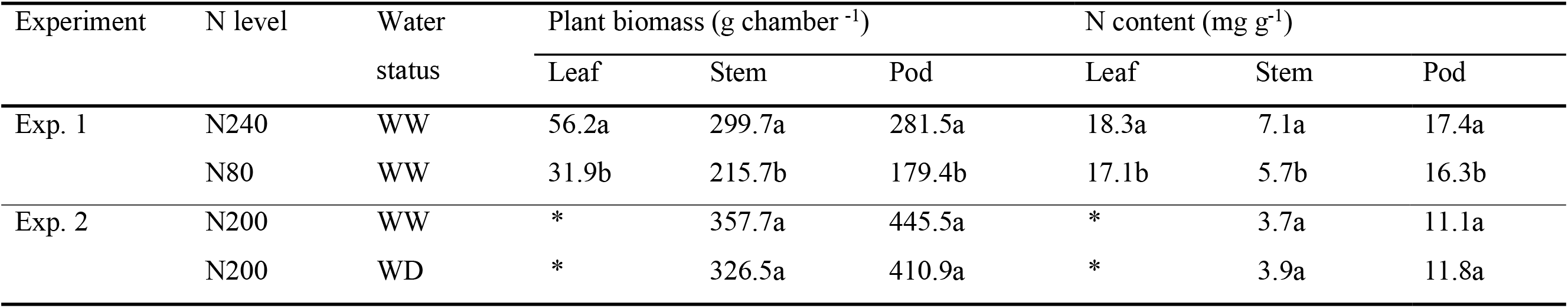
Mean plant dry matter and N contents for different N or water treatments at final collection (n = 6). In Exp. 1 the oilseed rape plants were grown under 240 kg N ha^-1^ (N240) or 80 kg N ha^-1^ (N80) applications and were all well-watered (WW); in Exp. 2 all plants were fertilized with 200 kg N ha ^-1^, and grew under well-watered or water deficit (WD) conditions. In the same column, different letters indicate significant differences between treatments in Exp. 1 (*p* < 0.05). No significant differences were found between water treatments in Exp. 2. Pods were not fully ripe in either experiments. *There were no leaves left when oilseed rape plants were harvested in 2020.

| Experiment | N level | Water status | Plant biomass (g chamber<sup>-1</sup>) |  |  | N content (mg g<sup>-1</sup>) |  |  |
| --- | --- | --- | --- | --- | --- | --- | --- | --- |
|  |  |  | Leaf | Stem | Pod | Leaf | Stem | Pod |
| Exp. 1 | N240 | WW | 56.2a | 299.7a | 281.5a | 18.3a | 7.1a | 17.4a |
|  | N80 | WW | 31.9b | 215.7b | 179.4b | 17.1b | 5.7b | 16.3b |
| Exp. 2 | N200 | WW | * | 357.7a | 445.5a | * | 3.7a | 11.1a |
|  | N200 | WD | * | 326.5a | 410.9a | * | 3.9a | 11.8a |

No significant differences were found in biomass or N content between the water treatments in Exp. 2. Plants that grew under lower soil water content tended to have a lower stem and pod biomass, while the N content in the pod and stem samples at harvest was slightly higher when less water was supplied, although not significant (Table 3).

### Root growth

Root growth was recorded from March to June, covering tracer injection and sampling periods in both years (Fig. 1). Roots were present below 1.7 m already in April in both experiments. In Exp. 1, roots reached just below 2 m depth during the labelling period (Fig. 1a). In Exp. 2 roots were present below 3 m in April (Fig. 1b). At the time of labelling, the average root intensities in the top 2 m soil layers were approximately four times higher in Exp. 2 than in Exp. 1 (0.25 and 0.06 cm cm^-2^, respectively).

**Fig. 1.**
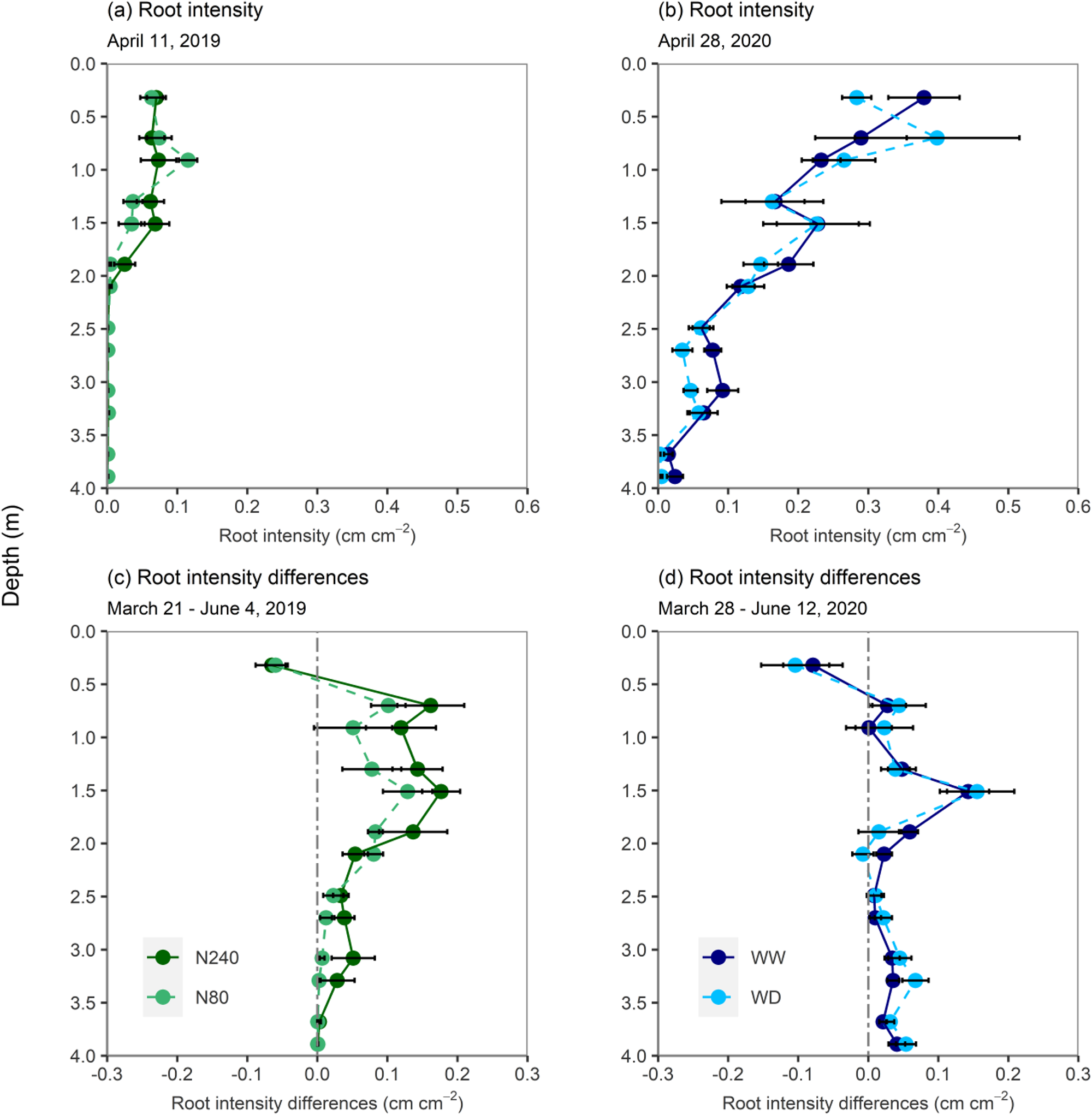
Root intensity measured on April 11, 2019 (a), 8 days after tracer injection and on April 28, 2020 (b), 11 days after tracer injection. Differences in root intensity from March 21 to June 4, 2019 (c) and from March 28 to June 12, 2020 (d). N240 = 240 kg N ha^−1^, N80 = 80 kg N ha^−1^, WW = well-watered, WD = water deficit. Error bars denote standard errors. No significant differences were found at any depths.

In both years and all treatments, root intensity tended to increase below 0.5 m from fertilization in March to June (Fig. 1c and d). There was a tendency towards more root growth in the N240 than in the N80 treatment in the lower soil layers. No significant differences in root growth were found between the two water regimes. In both experiments, the root intensity in the top 0.5 m decreased from March to June.

### Water extraction

VWC at the three recorded depths were similar in the two N treatments during the isotope sampling period in Exp. 1. Only the VWC at 0.5 m depth in the water deficit treatment tended to be lower during the sampling period in Exp. 2 (Fig. 2a and c).

**Fig. 2.**
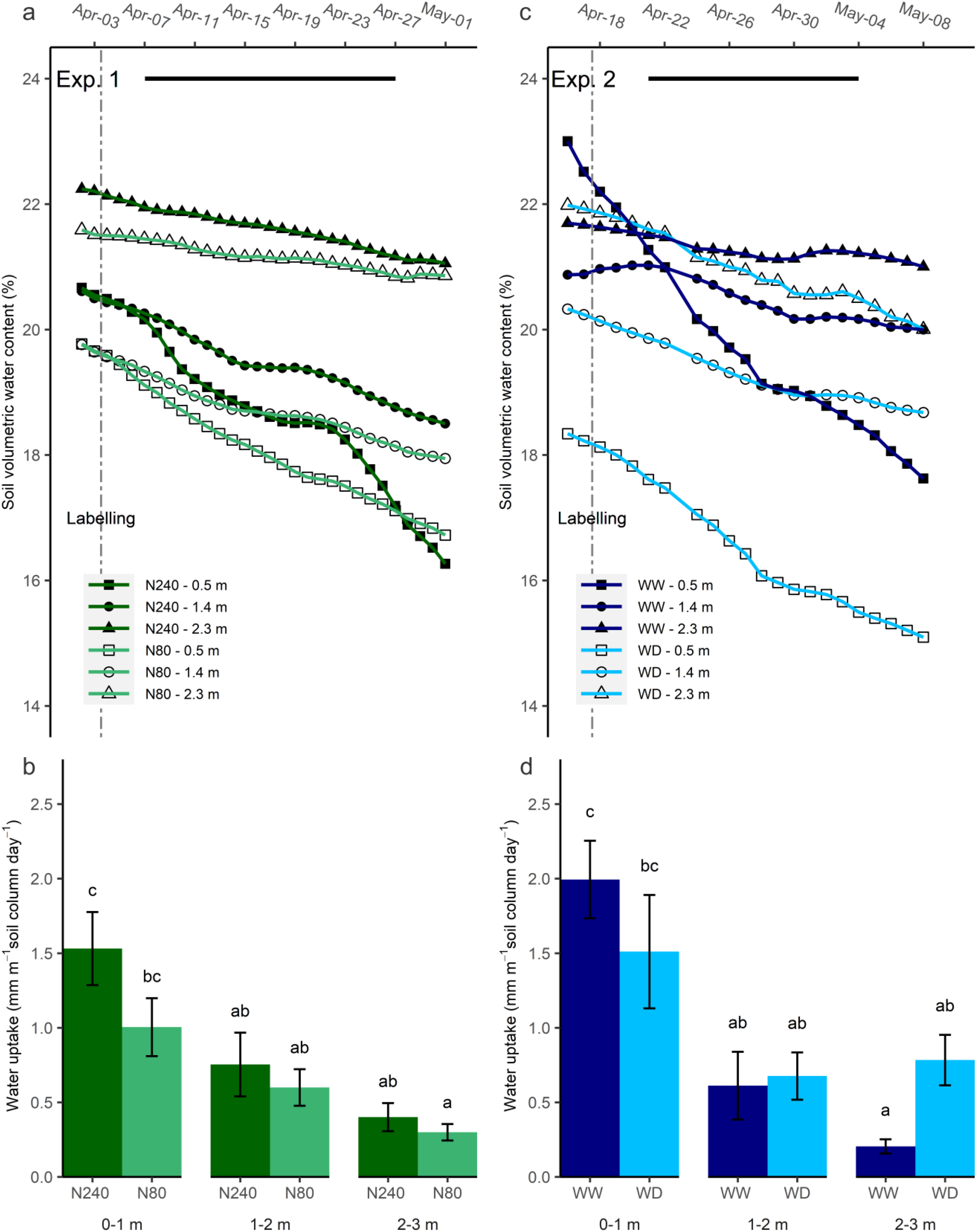
Soil volumetric water content (VWC; %) and water uptake in Exp. 1 (a, b) and Exp. 2 (c, d). N240 = 240 kg N ha^−1^, N80 = 80 kg N ha^−1^, WW = well-watered, WD = water deficit. Data were collected from April 2 to May 1, 2019 in Exp. 1 and from April 16 to May 8, 2020 in Exp. 2. Daily averages of recorded VWC are shown in a and c. The black segments denoted selected periods for daily water decrease estimations in Exp. 1 (b) and Exp. 2 (d). Daily water uptake from each 1 m interval of soil column was estimated as averages of daily water decrease from that column from April 7 to April 27, 2019 in Exp. 1 and from April 21 to May 4, 2020 in Exp. 2. Error bars denote standard errors, letters indicate significant differences across the treatments in the same experiment (p<0.05).

Based on the simplified estimations of daily water uptake, more than 1 mm of water was removed from the 0-1 m soil layer per day during the selected labelling period, while less than 1 mm was removed from the 1-2 and 2-3 m soil layers in Exp. 1 (Fig. 2b). It was clear that with higher N application, water uptake throughout the whole soil profile was increased, though not significant.

The total amount of water taken up in the two water regimes in Exp. 2 was similar. In total, 2.80 and 2.97 mm water per day was removed within the selected period from the top 3 m of the soil column in the WW and WD treatment respectively (Fig. 2d). However, a shift towards water uptake from deeper soil layers in the WD treatment was observed, as the water deficit in the topsoil increased water uptake in the 2-3 m interval. However, the trend was not significant. Additionally, slight and insignificant increases in δ^13^C values were observed in leaves, which further indicated plants under the WD treatment were not drought-stressed during the labelling period in Exp. 2 (Fig. 3).

**Fig. 3.**
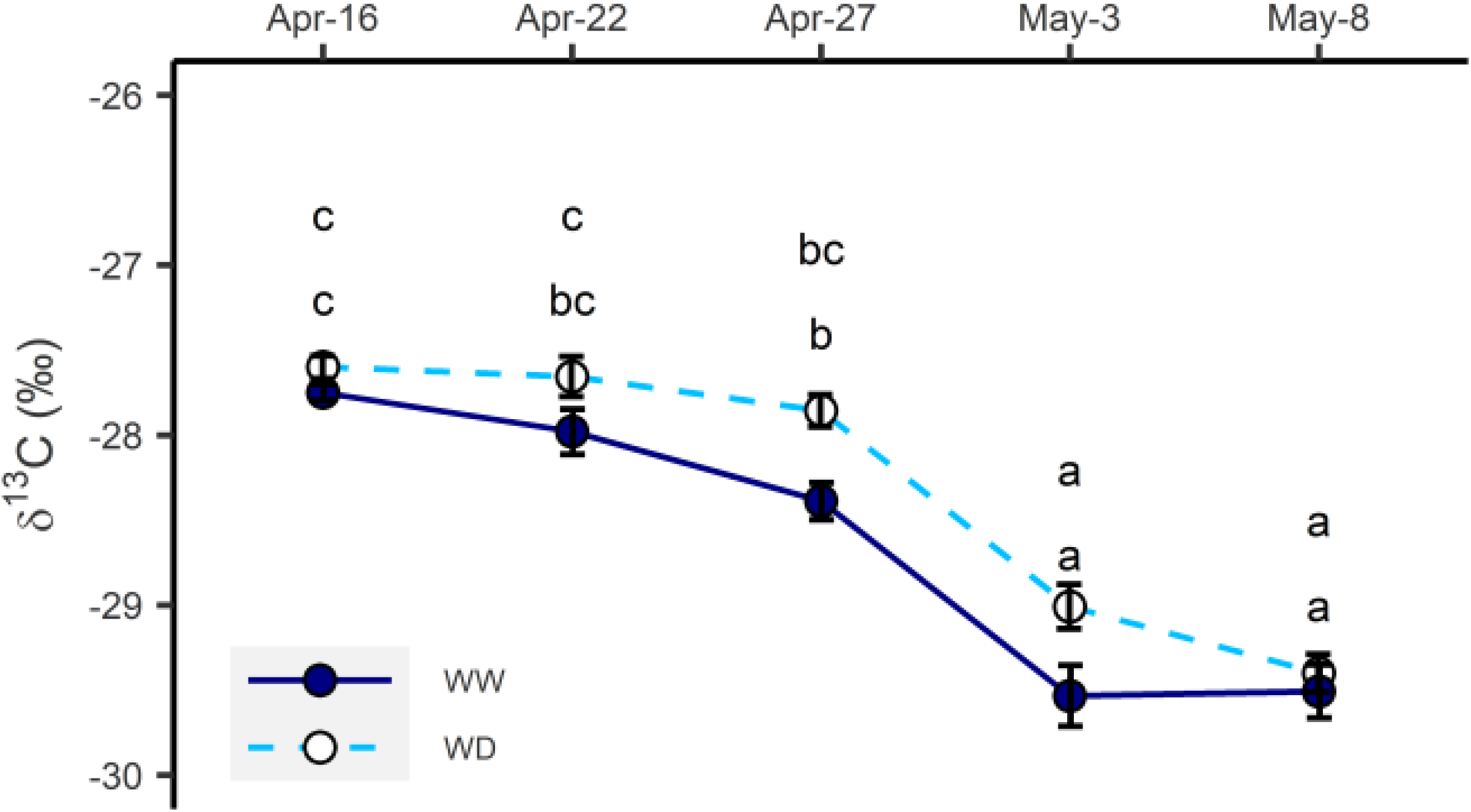
δ^13^C measured in leaf samples collected during the isotope sampling period in Exp. 2 in well-watered (WW) and water deficit (WD) treatments. Error bars denote standard errors, letters indicate significant differences across the treatments (p<0.05).

### ^2^H enrichment

At both N treatments, when the tracer was injected at 0.5 m depth instead of 1.7 m, a higher enrichment of ^2^H in transpiration water was expected (Fig. 4a). Besides, the ^2^H enrichment of the transpiration water was higher in N240 than in N80 treatments on all dates and both injection depths. The concentration of ^2^H in transpiration water increased significantly with time when the injection was conducted at 1.7 m. However, when ^2^H was injected at 0.5 m, no increase in concentration with time was observed.

**Fig. 4.**
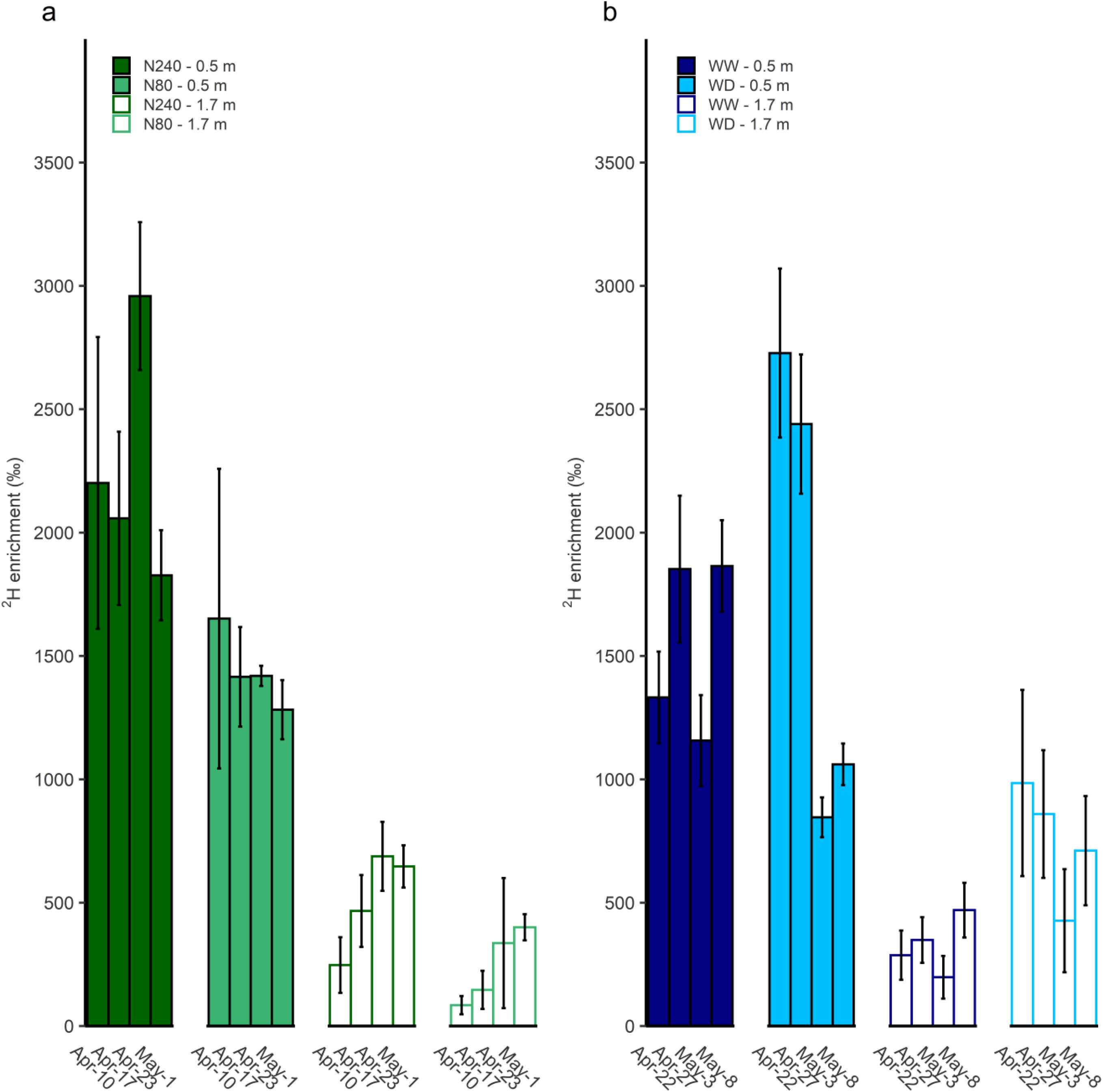
Time course of ^2^H enrichment in transpiration water was shown under N (a) or water (b) treatments during isotope sampling periods. N240 = 240 kg N ha^−1^, N80 = 80 kg N ha^−1^ (a), WW = well-watered, WD = water deficit (b). ^2^H labelled water was injected at either 0.5 or 1.7 m in each treatment. Error bars denote standard errors (n=3). Mean values are shown here (± SE).

During the labelling period in Exp. 2, the lowest ^2^H concentration in the transpiration water was found in the WW treatment when the tracer was injected at 1.7 m. When injected at 0.5 m depth, higher ^2^H concentrations in transpiration water at the WD treatment than at the WW treatment was seen at the first sampling dates, but in the WD treatment, it fell by c. 60% between April 27 and May 3, while it did not change much over time in the WW treatment (Fig. 4b).

In Exp. 1 the enrichments of ^2^H derived from 1.7 m were 5 – 35% of the ^2^H derived from 0.5 m in both treatments, showing a larger proportion of ^2^H uptake from 0.5 m than 1.7 m (Table 4). This ratio of deep and shallow derived ^2^H enrichment was increased 0 – 10 percentage points in the N240 than the N80 treatment during the sampling period. For Exp. 2, in the WW treatment, the ratio of ^2^H enrichment with tracer injected at 1.7 and 0.5 m was approximately 20% three weeks after injection. While in the WD treatment, the ratio was over 30% and further increased 30 percentage points at the last sampling date (Table 4).

**Table 4.**
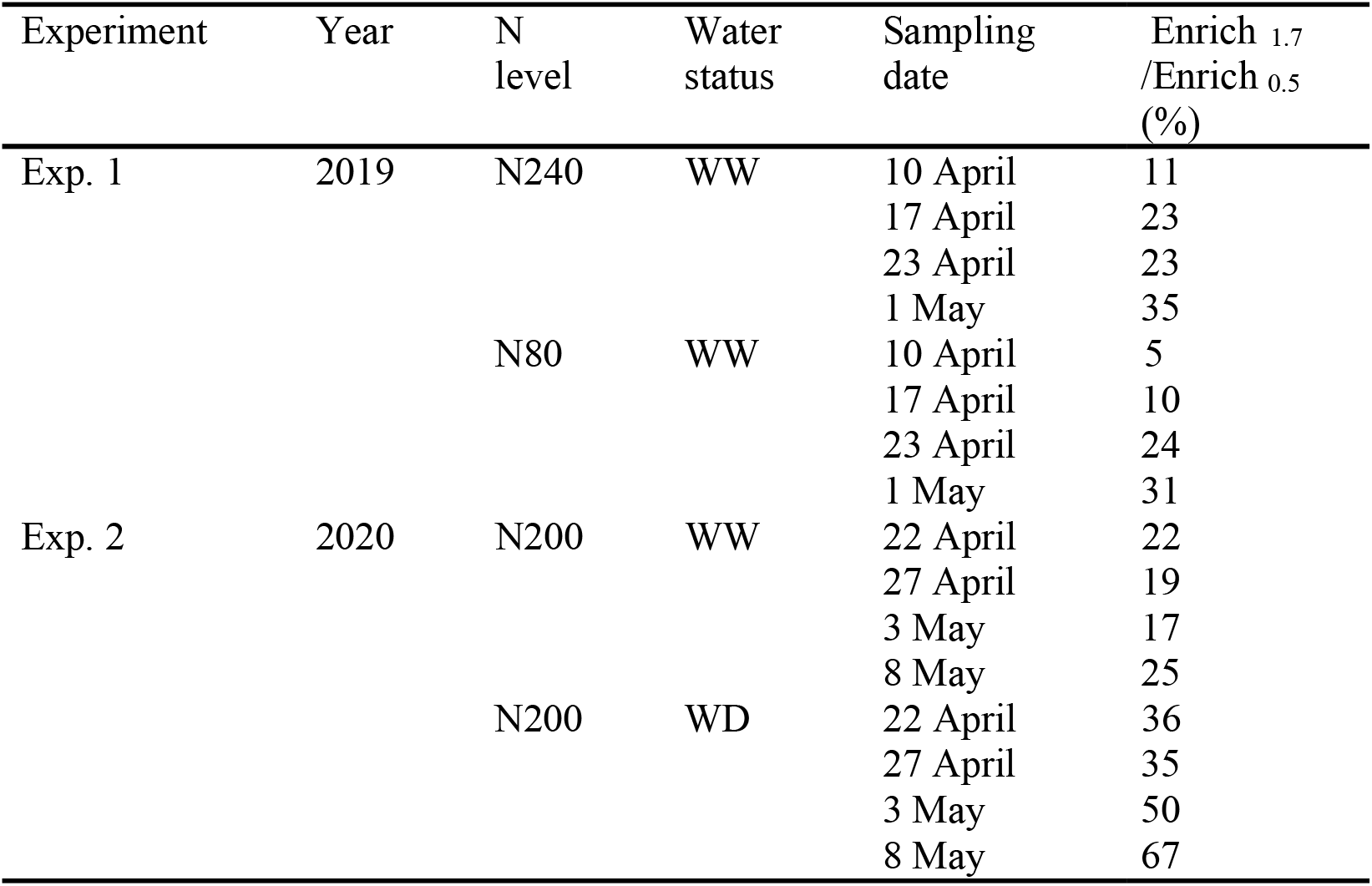
The ratio of ^2^H – enrichment in transpiration water with tracer injected at 1.7 m (Enrich _1.7_) and 0.5 m (Enrich _0.5_). The oilseed rape plants were grown under 240 kg N ha^-1^ (N240) or 80 kg N ha^-1^ (N80) applications and were all well-watered (WW) in Exp. 1; in Exp. 2 all plants were fertilized with 200 kg N ha ^-1^ fertilizer, and grew under well-watered or water deficit (WD) conditions.

| Experiment | Year | N level | Water status | Sampling date | Enrich<sub>1.7</sub> / Enrich<sub>0.5</sub> (%) |
| --- | --- | --- | --- | --- | --- |
| Exp. 1 | 2019 | N240 | WW | 10 April | 11 |
|  |  |  |  | 17 April | 23 |
|  |  |  |  | 23 April | 23 |
|  |  |  |  | 1 May | 35 |
|  |  | N80 | WW | 10 April | 5 |
|  |  |  |  | 17 April | 10 |
|  |  |  |  | 23 April | 24 |
|  |  |  |  | 1 May | 31 |
| Exp. 2 | 2020 | N200 | WW | 22 April | 22 |
|  |  |  |  | 27 April | 19 |
|  |  |  |  | 3 May | 17 |
|  |  |  |  | 8 May | 25 |
|  |  | N200 | WD | 22 April | 36 |
|  |  |  |  | 27 April | 35 |
|  |  |  |  | 3 May | 50 |
|  |  |  |  | 8 May | 67 |

### N depletion and accumulation

In spring both years, soil nitrate concentrations of the top 1.7 m soil were low (Table 5) and did not change much between the first and second sampling dates. There was a high variation in ^15^N concentration in the soil nitrate at the soil sampling after the isotope sampling period, and no significant differences were found between nitrate and ^15^N under different N treatments at any of the sampled depths. In Exp. 2, no significant differences were found in nitrate depletion between the WW and WD treatments (Table 5).

**Table 5.** Means of soil nitrate content and ^15^N concentration in different soil layers before and at the end of labelling periods (in 0.5 and 1.7 m, n=3; in 1.1 m, n=6). The oilseed rape plants were grown under 240 kg N ha^-1^ (N240) or 80 kg N ha^-1^ (N80) applications and were all well-watered (WW) in Exp. 1; in Exp. 2 all plants were fertilized with 200 kg N ha ^-1^ fertilizer and grew under well-watered or water deficient (WD) conditions. In Exp. 1, soil samples were taken on April 3 and May 3, 2019; in Exp. 2, soil samples were taken on April 17 and May 9, 2020. For soil nitrate content analysis, soil samples, which were taken from labelled depths, were excluded to avoid the effects of tracer application. For soil ^15^N concentration analysis, soil samples of 0.5 and 1.7 m were taken from labelled depths, soil samples of 1.1 m were taken from all depths.

| Experiment | N level | Water status | Depth (m) | Soil NO<sub>3</sub><sup>-</sup> before (mg N kg<sup>-1</sup> dry weight) | Soil NO<sub>3</sub><sup>-</sup> after (mg N Kg<sup>-1</sup> dry weight) | Soil $^{15}\text{N}$ before (‰) | Soil $^{15}\text{N}$ after (‰) |
| --- | --- | --- | --- | --- | --- | --- | --- |
| Exp. 1 | N240 | WW | 0.5 | 2.6 | 2.0 | 102 | 1116 |
|  |  |  | 1.1 | 2.4 | 2.2 | 19 | 33 |
|  |  |  | 1.7 | 2.9 | 2.2 | 163 | 8669 |
|  | N80 | WW | 0.5 | 2.4 | 1.7 | 364 | 1323 |
|  |  |  | 1.1 | 1.8 | 2.4 | 31 | 38 |
|  |  |  | 1.7 | 2.9 | 2.9 | 70 | 5109 |
| Exp. 2 | N200 | WW | 0.5 | 0.1 | 0.1 | * | * |
|  |  |  | 1.1 | 0.1 | 0.2 | * | * |
|  |  |  | 1.7 | 0.1 | 0.2 | * | * |
|  | N200 | WD | 0.5 | 0.1 | 0.1 | * | * |
|  |  |  | 1.1 | 0.1 | 0.1 | * | * |
|  |  |  | 1.7 | 0.1 | 0.1 | * | * |

\* Soil $^{15}\text{N}$ concentration was too low to detect.

Corrected for ^15^N already in the soil before tracer injection, additional ^15^N tracer resulted in higher ^15^N enrichment in the biomass samples (Fig. 5). At the cessation of Exp. 1, plants under the same N treatment, with ^15^N tracers injected at either 0.5 or 1.7 m, exhibited similar ^15^N uptake efficiency. ^15^N use efficiency of oilseed rape plants in the N80 treatment was twice as high as in the N240 treatment. In general, an extra gram of ^15^N led to a 12-15% increase in biomass ^15^N atom fraction in the N80 treatment, while in the N240 treatment, the increase in ^15^N atom fraction per gram ^15^N added was only around 6% (Fig. 5a). During the labelling period in Exp. 1, the leaf ^15^N uptake efficiency tended to increase with time (Fig. 5b), which indicated more ^15^N was accumulated in the leaves. However, this was only significant when ^15^N was injected at 1.7 m. Independently of ^15^N injection depths a larger fraction of the applied ^15^N was found in leaves of oilseed rape plants that had been fertilized with a lower amount of N.

**Fig. 5.**
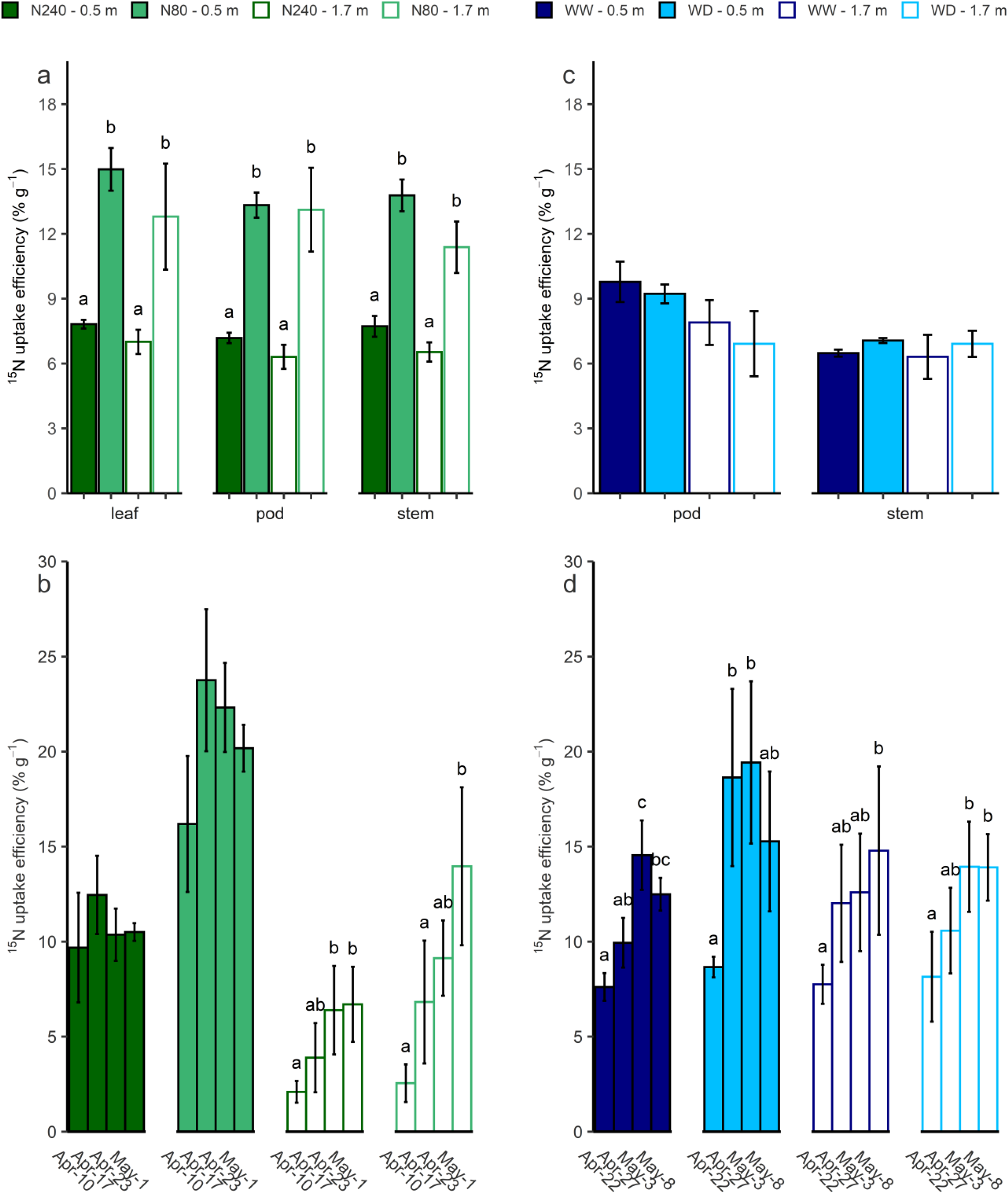
^15^N uptake efficiency (% g^-1^) that was measured in harvest samples (a, c), and leaf samples (b, d) which were collected during the labelling periods in Exp. 1 (a, b) where, N240 = 240 kg N ha^−1^, N80 = 80 kg N ha^−1^ and Exp. 2 (c, d) where, WW = well-watered, WD = water deficit. ^15^N tracer was injected at either 0.5 or 1.7 m in each treatment. Mean values of harvest ^15^N use efficiency in different organs under different N (a) or water (b) treatments are shown here (± SE). Error bars denote standard errors (n=3), and letters indicate significant differences among treatments within the same organ (p<0.05). Time course of leaf ^15^N use efficiency (% g ^-1^) under different N (c) and water (d) regimes was calculated. Error bars denote standard errors (n=3), and letters indicate significant differences among all sampling dates under the same treatment and injection depth (p<0.05). Mean values are shown here (± SE).

^15^N uptake efficiency was almost unaffected by the soil water status or injection depth in the harvest samples of Exp. 2 (Fig. 5c). While the fraction of ^15^N in leaves significantly increased with time, no clear water treatment or injection depth effect was observed at any measurement date (Fig. 5d).

## Discussion

### Effect of N and water supply on root growth

The difference in root growth along the soil profile between two N treatments was small in the current study. However, we found a non-significant tendency towards deeper and more roots in the subsoil after the high rate of N application. The effects of N supply on root growth were found to be inconsistent in previous studies. Svoboda and Haberle (2006) claimed that a high N fertilization rate led to reduced wheat rooting depth and density in deeper layers, while Hodge et al. (1999) found the increased soil N availability to lead to stronger shoot and root growth. The local soil N availability in subsoil was similar in the high and low N treatments. Thus, the observed enhancement of deeper root growth under high N supply was most likely a concurrent effect with better shoot growth.

According to previous studies, water deficiency has been proven to be the stimulator of deep root growth and the uptake of deep soil water (Bloom et al. 1985; Vandoorne et al. 2012). Surprisingly, the effect of water supply on root growth was not seen in this study. In both treatments, the increment of root intensity along the soil profile except at 1.5 m depth was less than 0.1 cm cm^-2^, indicating little and non-preferential root growth under short-term water deficit. One possible explanation for this could be that the water deficiency we observed in the experiment was not severe or long enough to stimulate the deep root growth previously observed (Skinner 2008; Vandoorne et al. 2012). The other explanation is that plants grew under the water deficit treatment were available to obtain adequate water from the subsoil. Therefore their growth was not affected by the water deficit.

Compared with roots in topsoil, the contribution of deep roots to resource uptake is often ignored, as they usually develop late and have a limited active period. However, deeper and denser roots in subsoil do indicate a better capacity for resource acquisition from deep soil layers (Kell 2011; Maeght et al. 2013). Thus, the factors that directly or indirectly affect deep root growth may also affect deep resource uptake.

### Effect of N and water supply on water uptake

N supply affected the amount, distribution, and dynamics of water uptake. With a higher N fertilization rate, root water uptake was higher at all depths along the soil profile, with most of the water taken from the top 1 m layer. Higher ^2^H enrichment in transpiration water was found in N240 treatment compared to N80 treatment wherever the tracer was applied. This could be an indicator for higher uptake of ^2^H-labelled water from the labelled depth, or showing that ^2^H-labelled water uptake from the other depths was lower (Rasmussen and Kulmatiski 2021). There is a possibility that the top-50-cm soil layer, where the soil VWC was not record, dried out during the experimental period, and topsoil in N240 treatment dried out faster than N80 treatment. In this case, plants fertilized with N240 took less water from the topsoil, leading to a higher ^2^H enrichment in transpiration water. However, as all plants were expected to be adequately supplied with water in the current experimental design, the higher ^2^H enrichment we found in N240 was mostly due to the higher water uptake from the labelled depth.

With higher N application, the water uptake is promoted by the increased aboveground growth, transpiration, and photosynthesis (Taylor et al. 1991; Waraich et al. 2011). In addition, high nitrate supply improves radial water fluxes in roots, as it up-regulates the expression of aquaporin and enhances root hydraulic conductivity (Gorska et al. 2008; Wang et al. 2016). Still, the improvement of water uptake was more evident at upper soil layers, where most of the roots were located and directly affected by the N supply.

In the second experiment, as the foliar δ^13^C values showed, water deficiency in this experiment did not seem to stress the oilseed rape plants, indicating plants under WD treatment took adequate water to maintain their growth. The relationships between ^13^C isotopic composition/discrimination and water stress have been widely studied for evaluating the performance of crop under water stress (Farquhar and Richards 1984; Farquhar et al. 1989; Dercon et al. 2006). Plant δ^13^C was reported to increase under water-limited conditions (Yousfi et al. 2012). Correspondingly, topsoil water status did not significantly affect the total amount of water uptake, while we observed altered water uptake distribution under reduced water supply. More water was taken from deep soil layers in WD treatment than WW treatment, which implied water deficit stimulated water uptake compensation from deep soil layers. This corresponds to previous findings by Vandoorne et al. (2012) and Hashemian et al. (2015), showing that moisture level of different soil layers is a key factor which affect root water uptake distribution. However, with a similar experimental setup, Rasmussen et al. (2020a) concluded that chicory failed in compensating water uptake from deeper soil layers. They suggested that the high hydraulic resistance and drought-induced stomatal closure might reduce root water uptake and plant water demand, leading to the failure of compensation (Rasmussen et al. 2020b).

Except in the WW treatment, the ratio of 1.7 m to 0.5 m derived ^2^H-enrichment increased with time. The increments were 26, 24, and 31 percentage points in N80, N240, and WD treatment, respectively. This distribution shift suggests the rising importance of deep roots in water uptake over the three-week period following injection. The effect of time on deep water uptake may be due to a direct effect on the development and maturity of roots, which has also been demonstrated by Garrigues et al. (2006), or to exhaustion of the labelled-^2^H at the 0.5 m depth.

### Effect of N and water supply on soil N content and N uptake

Our results showed that increasing N fertilization from 80 to 240 kg N ha^-1^ did not significantly change the content of inorganic N in the top 1.7 m soil, and water deficiency had little effect on the concentration and distribution of soil nitrate. However, these findings should be interpreted with caution as there were only a few weeks between the applications of N or water treatments until the soil measurements. In other studies, such treatments lasted longer, and oilseed rape grown under higher N fertilization rate or less water supply left more soil nitrate in topsoil layers (Smith et al. 1988; Dresbøll et al. 2016).

N uptake efficiency depends on both root uptake capacity and crop demand. Oilseed rape needs high N input during the vegetative growing stage (Rathke et al. 2006). Nevertheless, even with a high capacity to absorb N in autumn and winter, the recovery of fertilized N by oilseed rape is generally found to be poor (Sieling and Kage 2010), and an increasing rate of N supply can further reduce the N recovery (Rathke et al. 2006; Bouchet et al. 2016). This reduction in N recovery was also observed in our study. Despite the fact that the biomass in the N240 treatment was higher than the N80 treatment, ^15^N recovery in the biomass was higher in the N80 treatment, which indicated more thoroughly soil N depletion under a lower N fertilization rate. Svoboda and Haberle (2006) pointed that the effect of high nitrogen supply in the topsoil on reduction of N depletion in the subsoil can be the result of both less N demand from the subsoil, and reduced root growth in the subsoil. In our case, the latter was not observed.

Effects of water supply on ^15^N uptake efficiency were not observed in the current study. This contrasts with others’ findings, which demonstrated that water deficit would reduce N uptake and use efficiency in various ways (Jensen et al. 1997; Gonzalez-Dugo et al. 2010; Sadras et al. 2016; Riar et al. 2020). In general, topsoil water deficit reduces the availability of soil N (Jensen et al. 1997) and restricts its movement via mass flow and diffusion (Plett et al. 2020), hence reducing N uptake from top layers. To meet crop demand, the N uptake from subsoil would be stimulated. However, it does not seem to be the case in our study. Two possible explanations may account for the absent observation of compensated N uptake; one is that the extent of water deficit was not so severe that it affected soil N availability and subsequent N uptake. The other is that soil dryness directly reduced crop N demand via reducing the shoot growth; therefore, the efficient N uptake from subsoil is no longer needed. As no significant reduction in biomass was found in our case, we assume the second explanation was not the valid reason for the absence of enhanced deep N uptake.

The injection depth did not affect ^15^N uptake efficiency measured in harvest biomass samples in either experiment, while the continuous leaf sampling after injection showed different patterns of ^15^N uptake dynamic at 0.5 and 1.7 m. When ^15^N-labelled nitrate was applied at 0.5 m, the ^15^N uptake efficiency reached a peak in one to two weeks, while the efficiency kept increasing after injection at 1.7 m, suggesting some ^15^N-labelled nitrate still remained in the soil. Besides, in Exp. 1, one month after the injection, soil ^15^N enrichment at 1.7 m was much higher than at 0.5 m, which further confirmed that more ^15^N had been left at 1.7 m after injection. The continuous and delayed N uptake from subsoil was also observed in winter wheat by Haberle et al. (2006). They suggested the inadequate N supply from topsoil might be the stimulator of subsoil N uptake. Still, ^15^N uptake was less affected by deep resource placement than ^2^H uptake during the three-week sampling period. This indicates that while we see substantial uptake of both water and N by the deep parts of the root system, uptake of the two resources is not equally limited by the low root density and the short time available for active uptake (Chen et al. 2021).

There is no doubt that supplemental N and water supply also affect biomass production. In general, oilseed rape plants grown with extra N and water supply have been shown to have a higher yield (Taylor et al. 1991; Schjoerring et al. 1995; Dresbøll et al. 2016; Riar et al. 2020). In the current study, higher fertilization rate exhibited positive effects on biomass and plant N content. However, we only observed slight and non-significant increases of biomass under well-watered condition, together with a non-significant decrease in plant N content. This showed that the overall growth and development of oilseed rape plants were not restricted by water deficiency.

## Conclusions

Overall, when roots are already well developed in upper soil layers, N and water supply could still alter the water and N uptake via regulating deep water and N acquisition. The effects of reduced N and N supply on deep water and N uptake were not always the same. Increased water uptake from deep soil layers was not always accompanied by increased nitrogen uptake. Further research on the optimal water and N management strategies and the corresponding response of deep root functioning will be required to maximize the benefits of deep roots and maintain biomass production.

## Acknowledgements

We thank technician Aymeric d’Herouville for his contribution to the experimental work. We acknowledge the Villum Foundation (DeepFrontier project, grant number VKR023338) for financial support for this study, and China Scholarship Council for the financial support of Guanying Chen for her PhD research.

## Declarations

### Availability of data and material

The data supporting the findings of this study are available from the corresponding author, Guanying Chen, upon request.

### Code availability

Not applicable

### Authors’ contributions

Kristian Thorup-Kristensen, Dorte Bodin Dresbøll and Guanying Chen initiated the research, designed the experiment. Guanying Chen carried out the experiment and wrote the manuscript with support from Kristian Thorup-Kristensen and Dorte Bodin Dresbøll. Camilla Ruø Rasmussen helped with data analysis and participated in manuscript revision. Abraham George Smith provided the technical assistance in processing root images with software and offered language editing.

